# Drought-induced mortality: stem diameter variation reveals a point of no return in lavender species

**DOI:** 10.1101/848879

**Authors:** Lia Lamacque, Guillaume Charrier, Fernanda dos Santos Farnese, Benjamin Lemaire, Thierry Améglio, Stéphane Herbette

## Abstract

In the context of climate changes, water availability is expected to severely decline. Consequently, there is a need to predict mortality of woody species, especially to find a physiological threshold to drought-induced mortality. Lavender species (*Lavandula angustifolia* and *Lavandula x intermedia*) which are important crops of the Mediterranean region are affected by a decline, notably caused by successive intense drought events. Lavender response to extreme drought events was monitored using continuous stem diameter measurements. Water potential, stomatal conductance, loss of xylem hydraulic conductivity and electrolyte leakage were also measured during desiccation, and recovery was evaluated after rewatering. Two parameters computed from stem diameter variations were related to stress intensity and resilience to stress: PLD (Percentage Loss of Diameter) and stem PLRC (Percentage Loss of Rehydration Capacity of the stem), respectively. We showed that plants did not recover when the PLD reached its maximal value (PLD_max_) which was 21.27 ± 0.57% in both lavender species and whatever the growing conditions. This point of no return was associated with a high level of cell lysis evaluated by electrolyte leakage, and occurred far after the xylem hydraulic failure. We discussed the relevance of PLD_max_ as a threshold for drought-induced mortality and its physiological significance, in relation to the mortality mechanisms.

**One-sentence summary:** Under extreme drought, lavender death occurs when the water storage of the elastic compartment of the stem is exhausted.

## Introduction

Global climate change is inducing increasing temperature and precipitation changes resulting in longer and more severe drought events. Extreme drought events affect many agroecosystems worldwide (Allen et al., 2010). The Mediterranean region is one of the most impacted in terms of severe water-stress with high reduction in precipitation (Thuiller et al., 2005). In this area, the xerophilous shrubs lavender (*Lavandula angustifolia* Miller) and lavandin (*Lavandula x intermedia*) are crop species that are facing a major decline involving pathogen attacks coupled to a sharp reduction in rainfalls (Chuche et al., 2018; Sémétey et al., 2018). Although the maintaining hydraulic functioning has been reported as critical for plants survival facing drought stress (Brodribb and Cochard, 2009), the physiological thresholds leading to plants death are still fuzzy.

According to the tension-cohesion theory (Dixon and Joly, 1895), water flows under tension in the xylem conduits. The stomata close to avoid a high loss of water and so an increase in the xylem tension (Brodribb and Holbrook, 2003). However, the water potential continues to decrease and water is still lost through cuticular conductance, stomatal leakiness or bark (Choat et al., 2018). Tensions increase in the conduits and water columns can thus break by cavitation, a physical phenomenon by which a liquid phase is suddenly transformed into a gas. Xylem conduits are filled by air bubbles, creating embolism which blocks water transport and would lead to organ death and then plant death (Brodribb and Cochard, 2009; Barigah et al., 2013b). The lethal water potential in most of angiosperm trees is close to the one inducing high level of embolism, such as *P*_88_ (water potential inducing 88% loss of hydraulic conductance, Barigah et al., 2013b; Urli et al., 2013) or even *P*_99_ (Li et al., 2015). However, in conifers and grapevine, the lethal water potential has been defined close to the *P*_50_ (Brodribb and Cochard, 2009; Charrier et al., 2018). The theoretical lethal water potential has been observed and exceeded in still living angiosperm or conifer trees (Nardini et al., 2013; Adams et al., 2017; Hammond et al., 2019). Moreover, artefacts related to hydraulic measurements has been reported, in particular in case of high tension (at midday or under water stress, Wheeler et al., 2013). In order to improve the understanding of woody species response to severe drought, it is necessary to identify accurate physiological indicators for death, *i.e*. physiological thresholds that, if exceeded, would affect their ability to recover from a severe drought and in particular using a non-invasive sensor.

Water limitation affects growth as well, and the stem diameter variation can be a good indicator of mild water stress (Fernández and Cuevas, 2010). The variation in stem diameter exhibits a diurnal rhythm: contracting during daytime and expanding during nighttime (Offenthaler et al., 2001). Daily variations are related to the transpiration process and consequently to climatic variables such as air temperature, relative humidity, rainfalls and VPD (Devine and Harrington, 2011). Daily variation is a convolution of four independent components: i) irreversible radial growth, ii) reversible living-cell dehydration-rehydration, iii) thermal expansion and contraction and iv) expansion of dead conducting elements due to the increase and relaxation of internal tensions (Daudet et al., 2004). Dendrometers with continuous measurements allow monitoring tree water status and growth with several advantages: non-destructive, real time and automatic recording, high precision (Huck and Klepper, 1977; Ameglio and Cruiziat, 1992; Ortuño et al., 2010). The response in stem diameter under severe drought stress has rarely been investigated. Under mild water stress, the diameter variation includes secondary growth and changes in the water storage, so they are barely distinguishable (Daudet et al., 2004; Steppe et al., 2006). Under severe water stress, the diameter variation would only reflect fluctuations in the water storage as growth is inhibited.

In this study, we hypothesized that the loss in stem diameter would be a proxy of the water status and stress intensity and we tested its relevance as a threshold for drought-induced mortality. We have monitored stem diameter variations using LVDT dendrometer during mild and extreme drought events and following recovery in lavender plants grown under independent and different conditions. Two parameters computed from stem diameter variations were related to stress intensity and resilience to stress: PLD (Percentage Loss of Diameter) and stem PLRC (Percentage Loss of Rehydration Capacity of the stem), respectively. They were analyzed in relation to critical drought related traits including water potential, stomatal conductance, native embolism, vulnerability to embolism and electrolyte leakage to assess the cell lysis.

## Results

Well-watered (WW) and water-limited (WL) growing conditions exhibited significant differences in predawn water potential (*p* = 0.002 and <0.001 for (*Lavandula x intermedia* (*L.i.)* and *Lavandula angustifolia* (*L.a.)*, respectively) and maximal stomatal conductance (*p* <0.001 and 0.006 for *L.i.* and *L.a.*, respectively) before the first irrigation stop (IS_1_) (Table 1), highlighting acclimation to growing conditions in the experiment 2.

**Table 1.** Comparison of the two growth conditions wellwatered (WW) and water limited (WL) in 2018 in *L.a* and *L.i* for predawn water potential (*Ψ*_p_) and stomatal conductance (*g*_s_) measured just before the drought experiment. Means are significantly different between growth conditions according to Student Test at *p* < 0.05.

|  | Species | WW | WL | *p*-value |
| --- | --- | --- | --- | --- |
| $\Psi_p$ | *L.i* | $-0.104 \pm 0.007$ | $-0.260 \pm 0.04$ | 0.002 |
| | *L.a* | $-0.102 \pm 0.004$ | $-0.496 \pm 0.12$ | <0.0001 |
| $g_s$ | *L.i* | $348.85 \pm 19.64$ | $179.02 \pm 27.36$ | <0.0001 |
| | *L.a* | $258.55 \pm 15.88$ | $171.38 \pm 27.25$ | 0.0062 |

All the results obtained with the LVDT dendrometer during the dehydrations are schematized in Figure 1. Two parameters were computed: the percentage loss of diameter (PLD) and the percentage loss of rehydration capacity (PLRC).

**Figure 1.**
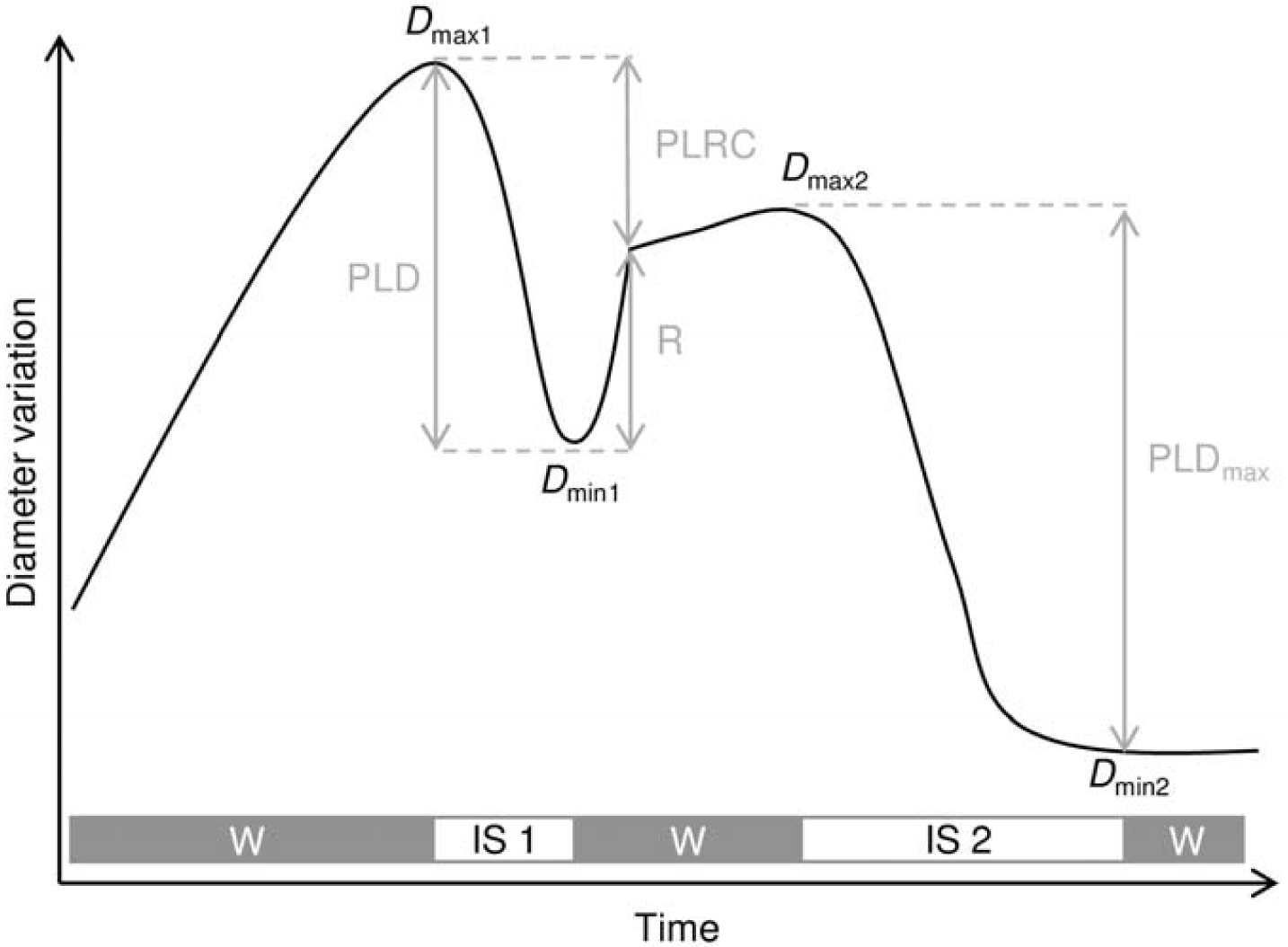
Schematic of the branch diameter variation versus time during the greenhouse experiments. The bar represents the watering conditions with watering (W) and irrigation stop (IS) in grey and white, respectively. *D*_max_ and *D*_min_ represent the maximum diameter before each IS and the minimum diameter before each rewatering, respectively. *D*_max1_ and *D*_min1_ were associated to the first IS and *D*_max2_ and *D*_min2_ were associated with the second IS. PLD corresponds to the percentage loss of diameter associated to an IS. R corresponds to the part of the diameter immediatly recovered following the rewatering. PLRC corresponds to the percentage loss of rehydration capacity, *i.e* the part of the diameter non recovered immediatly after a rewatering. When no recovery was observed after a rewatering, PLD_max_ and PLRC_100_ were reached.

Among the numerous dynamics of stem diameter variation (Supplemental Figure S1), typical patterns were observed depending on growing conditions and drought duration (Fig. 2). In all the recorded diameter variation dynamics, daily fluctuations were observed along the entire monitored period with a decrease and an increase in diameter during the day and night, respectively (Fig. 2). A phase of initial growth was observed during the initiation of floral buds and flowering followed by a slower growth. Water stress and rehydration periods exhibited rapid diameter change. The typical pattern shown in Figure 2A was only observed in WW plants (3 *L.a.* and 4 *L.i.*). During the first irrigation stop (IS_1_), the diameter exhibited a small decrease. During the second irrigation stop (IS_2_), the diameter decrease strongly while the water potential dropped below −9MPa after *ca.* 30 days. After rewatering, no recovery was observed neither in the short term (stem diameter recovery) nor in the longer term (regrowth the following year). The plants exhibiting this pattern were thus considered as dead and the computed PLD_max_ was associated to a PLRC value of 100. Daily fluctuations in branch diameters, although attenuated, remained in dead plants. However, the diameter exhibited a phase shift, increasing during the day and decreased during the night (Supplemental Figure S2). The typical pattern shown in Figure 2B was observed in both WW (5 *L.a.* and 5 *L.i.*) and WL plants (6 *L.a.* and 6 *L.i.*). During IS_1_, both diameter and midday water potential decreased to −2.8 ± 0.20 MPa and −2.55 ± 0.12 MPa for WW and WL respectively. After rewatering, the diameter quickly but partially recovered, whereas the water potential recovered (−1.6 ± 0.08 MPa and −0.72 ± 0.09 MPa for WW and WL, respectively). Then, the growth rapidly resumed and diameter *D*_max1_ was exceeded within a few days. After IS_2_, the water potential dropped below −9MPa and the diameter decreased and remained steady even after rehydration. All the plants exhibiting this pattern did not recover the following year and were therefore considered as dead. The typical pattern shown in Figure 2C was observed in WW plants only (3 *L.a.* and 2 *L.i.*). After IS_2_, the diameter and the water potential decreased (−5.86 ± 1.02 MPa). The second drought event was recoverable and growth resumed. The water potential of the individual exhibiting this pattern has promptly recovered (−0.76 ± 0.15 MPa).

**Figure 2.**
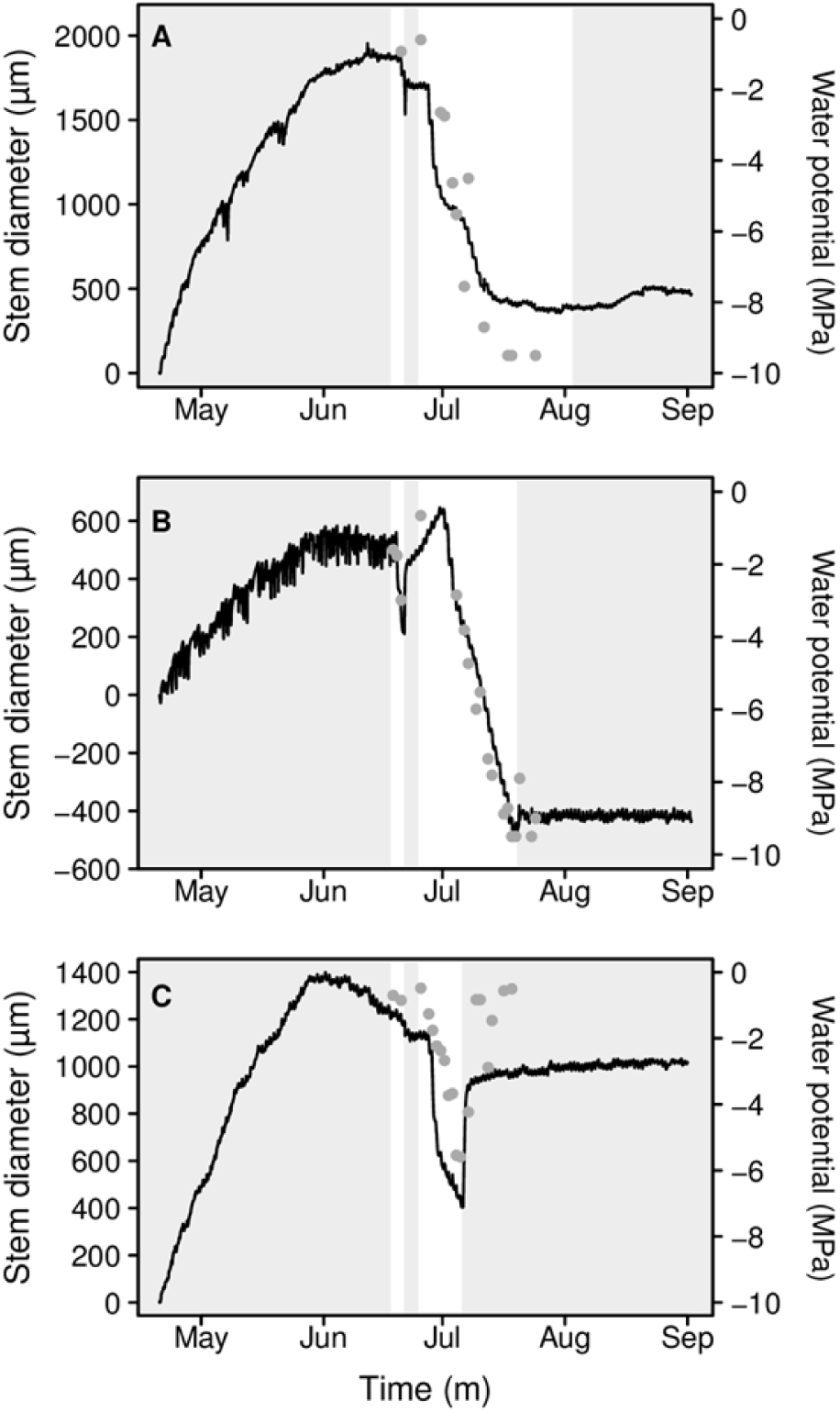
Representative example graphs illustrating 3 branch diameter variation dynamics (black line) and water potentials (grey points). A, *L.a.* in well watered (WW) condition with no recovery after the last irrigation stop (IS). B, *L.a.* in water limited (WL) condition with no recovery after the last IS. C, *L.a.* in WW condition with a recovery after the last IS. Shaded and white areas represent periods with and whitout watering, respectively. The diameter variation is represented from 0 at the beginning of the experiment.

PLD_max_ mean values were similar across growing conditions, species and experiments (*p* = 0.278, Table 2) and were not affected by the initial diameter of the branch (from 2,800µm to 9,200µm, R^2^ = - 0.019, *p* = 0.561). The mean PLD_max_ associated with lack of recovery was observed at 21.27 ± 0.57 %.

**Table 2.** Maximal loss of diameter (PLD_max_) scored for each experiment, growing condition and species (means ± se). Groups are not significantly different according to Kruskal-Wallis at *p* < 0.05.

| Year | Experimentation | Species | Condition | PLD<sub>max</sub> (%) |
| --- | --- | --- | --- | --- |
| 2017 | Greenhouse | *L.a* | WW | $20.84 \pm 1.48$ |
| | | *L.i* | WW | $21.24 \pm 1.40$ |
| 2018 | Greenhouse | *L.a* | WW | $21.96 \pm 0.42$ |
| | | *L.i* | WW | $23.49 \pm 2.39$ |
| | | *L.a* | WL | $20.48 \pm 1.46$ |
| | | *L.i* | WL | $18.37 \pm 1.26$ |
| | Temperature-controlled chamber | *L.a* | WW | $22.86 \pm 1.09$ |
| Mean PLD<sub>max</sub> (%) |  |  |  | **$21.27 \pm 0.57$** |

The ratio of xylem and bark were calculated on cross sections of control and dry plants of the drought experiment 1 (Table 3). Cross sections were made at the location of friction-free core of dendrometers and were observed with a binocular loupe. In hydrated plants, bark represented 23.81% ± 0.032 in *L.a.* and 21.29% ± 0.009 in *L.i.*. In dried plants, bark represented 16.28% ± 0.016 in *L.a.* and 15.96% ± 0.006 in *L.i.*.

**Table 3.**
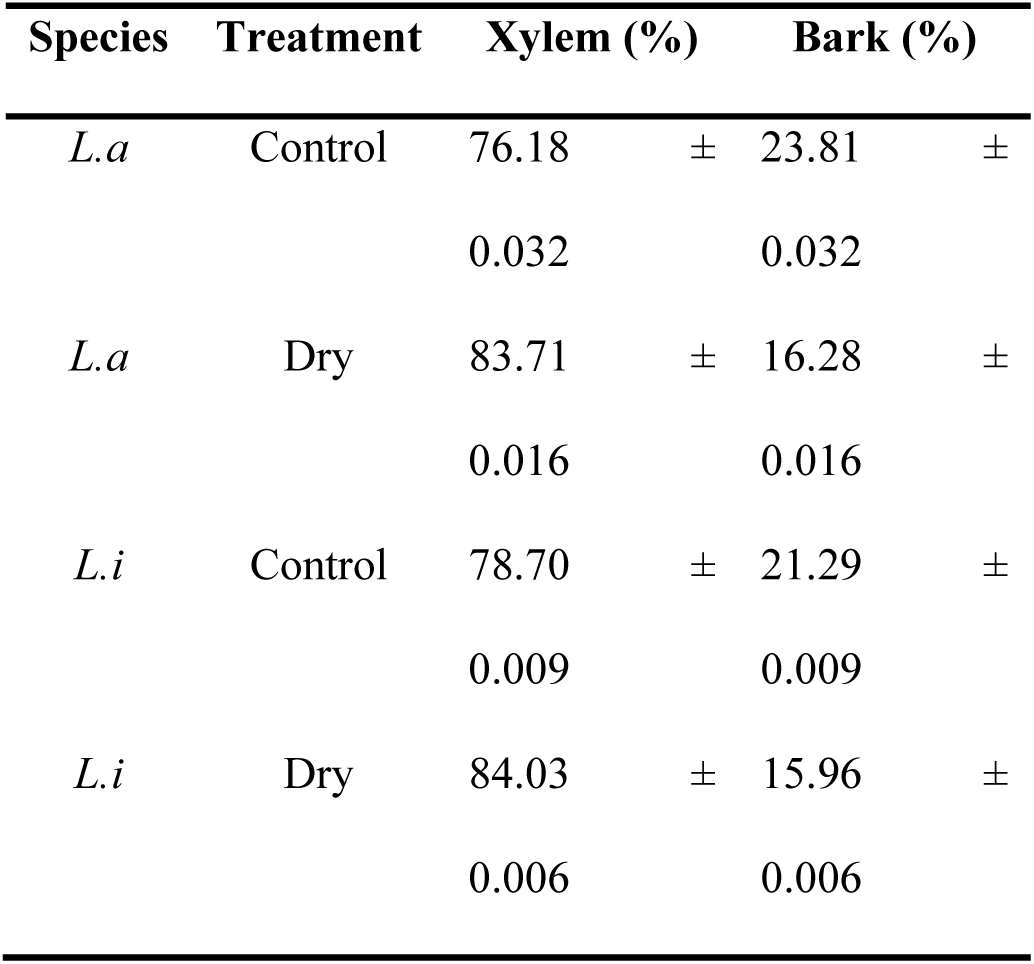
Ratio of xylem and bark on cross sections of Lavender species at the end of the drought experiment in 2017. Data are means ± standard error (n = 5).

PLD was positively correlated with PLRC (Fig. 3), in all species and growing conditions, according to a sigmoidal function (pseudo-r^2^ = 0.93 and 0.85 for *L.a.* and *L.i.*, respectively, Fig. 3). The PLD for which the PLRC reach 50 % (PLD_50_) was not significantly different across species (*p* = 0.273, 13.54 ± 0.39 % and 13.62 ± 0.58 % for *L.a.* and *L.i.*, respectively).

**Figure 3.**
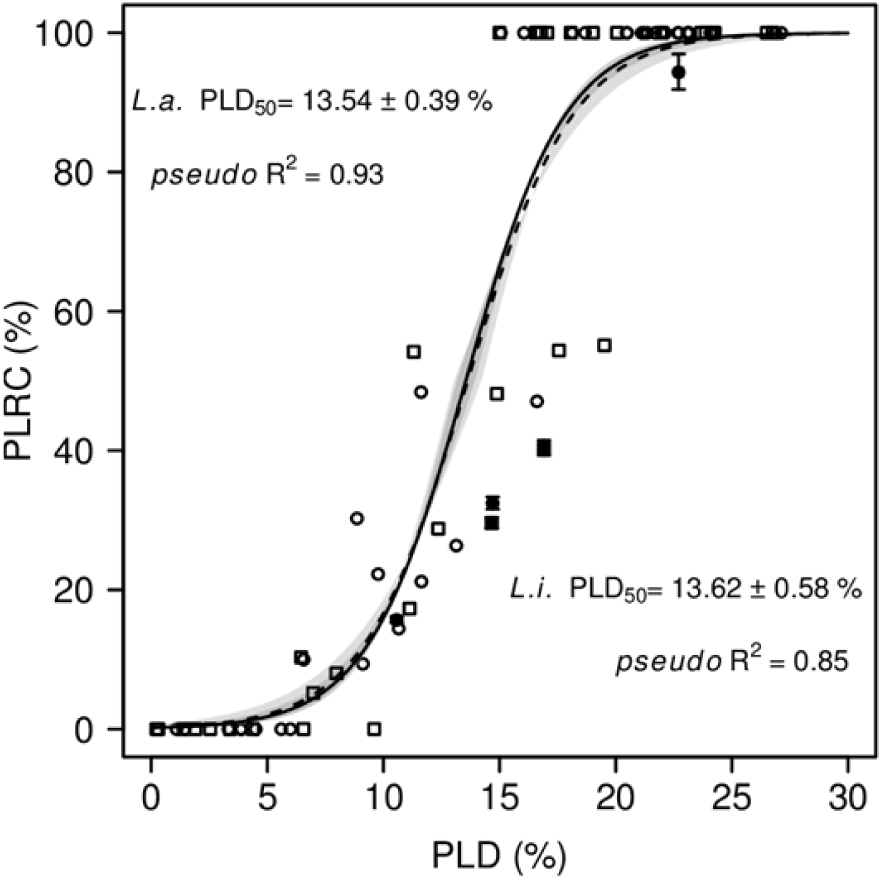
Stem Percentage Loss of Rehydration Capacity (PLRC) plotted versus Percentage Loss of Diameter (PLD) for *L.a.* (circles) and *L.i.* (squares). Data were obtained from 55 individual plants from the different conditions (experiment 1 and 2, well watered and water limited conditions, greenhouse and temperature-controlled chamber). For 5 individuals still alive, the PLD_max_ was not available, so their PLRC was estimated (black points) using mean PLD_max_ (Table 1). Dashed and dotted line are logistic-fit model and colored areas represent standard error for *L.a.* and *L.i.* respectively.

The *P*_50_ assessed by the Cavitron method were −2.46 ± 0.14 MPa and −2.41 ± 0.09 MPa for *L.a.* and *L.i.* respectively (Fig. 4B). Such a result means that these species are rather vulnerable to embolism. Therefore, another measurement of VC were performed by bench dehydration in experiment 1 (Fig. 4A) and it showed again high *P*_50,_ −2.86 ± 0.09 MPa and −2.71 ± 0.06 MPa for *L.i.* and *L.a.*, respectively. No significant differences were found between the two species (*p* = 0.748) and across the 2 methods (*p* = 0.767).

**Figure 4.**
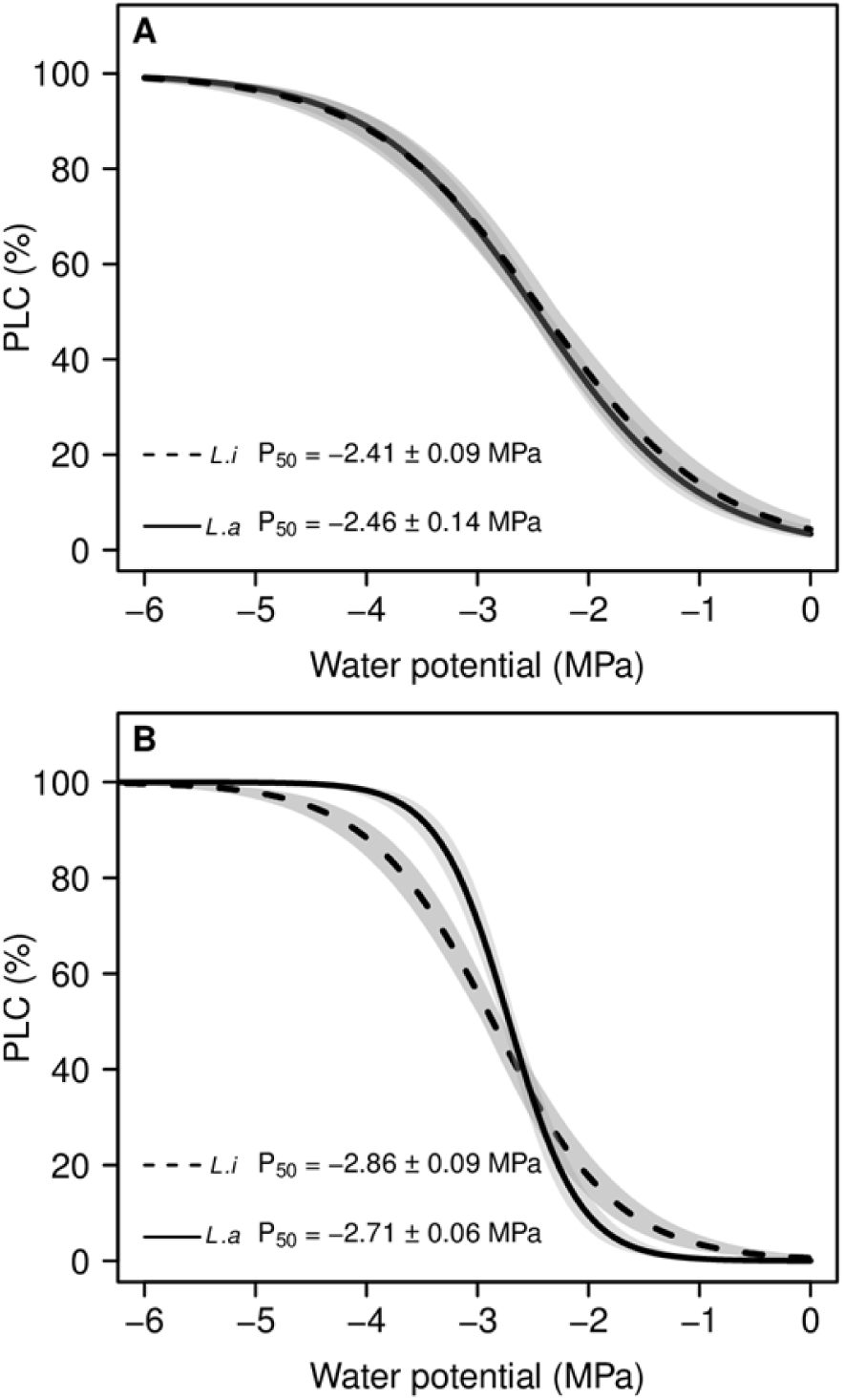
Vulnerability to embolism of *L.a.* (full line) and *L.i*. (dashed line). A, curves were obtained by the Cavitron method. B, curves were obtained by bench dehydration. Each curve is the mean curve per species from plants of the drought experiment 2 (n = 4 and 7 for *L.a.* and *L.i.*, respectively). Shaded areas are standard errors.

In both species, the water potential decreased nonlinearly with the PLD (Fig. 5, A and B). The water potential remained broadly stable up to 5% PLD and sharply decreased between 10 and 25% PLD. The stomatal conductance decreased with the PLD (Fig. 5A and B) and stomata were closed at near 5% PLD.

**Figure 5.**
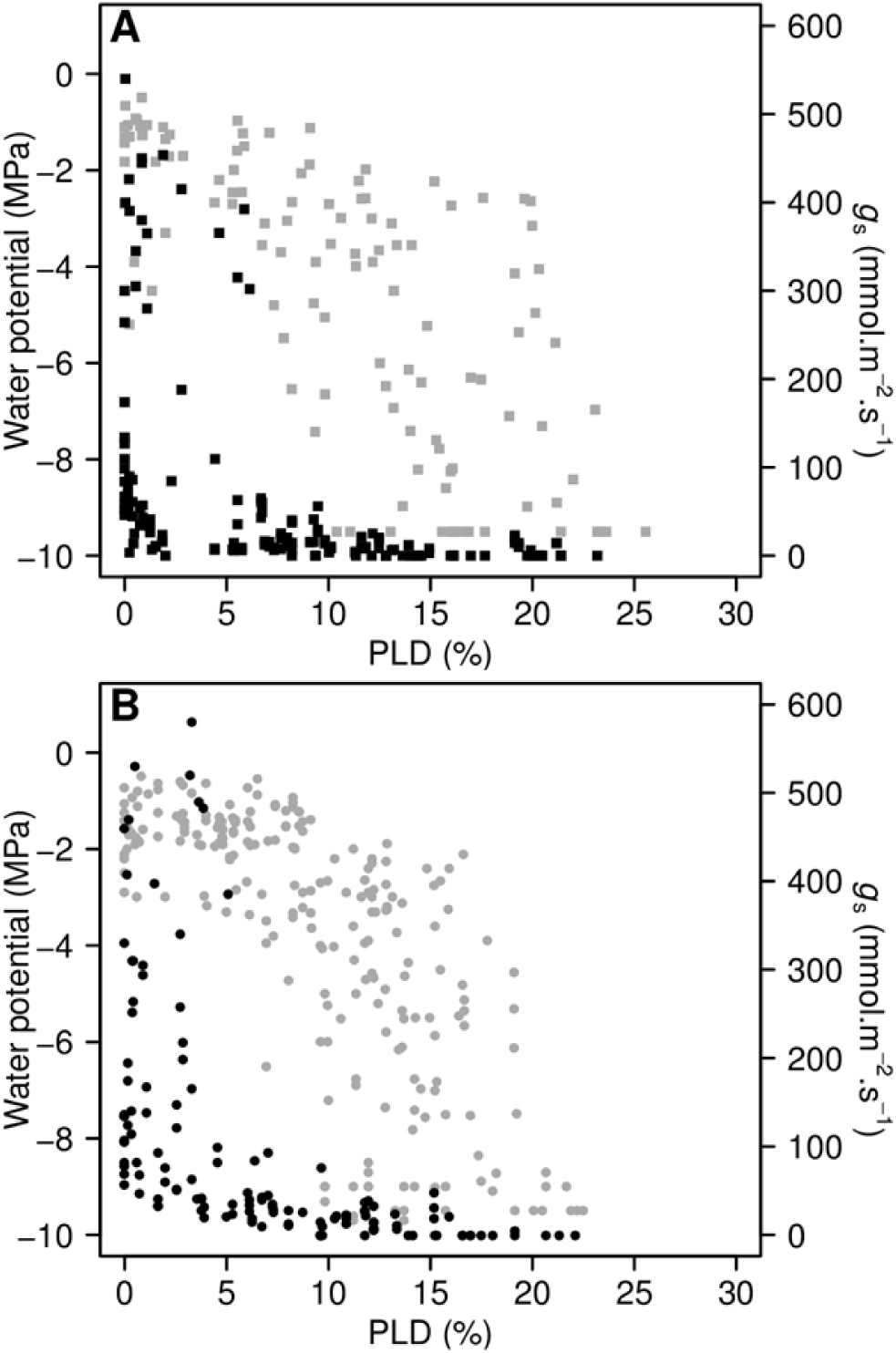
PLD-course of water potential (grey points) and stomatal conductance (black points) in *L.i.* (A) and *L.a.* (B). Data are individual measurements. At each point of the dehydration kinetics, at least 3 individuals per species and growth conditions were measured. PLD values were obtained from diameter variation measurements.

Under controlled conditions, at the beginning of the dehydration (0% PLD) in *L.a.*, PLC was at 9.46 ± 6.16 % and EL at 0 ± 2.30% (Fig. 6). At *ca.* 15% PLD, the PLC increased to 87.32 ± 3.61% (*p* = 0.0001, compared to the initial point) and EL increased to 47.22 ± 10.03 % (*p* = 0.003). At the end of the experimentation (*ca.* 25% PLD), PLC was at 97.37 ± 1.14 % (*p* = 0.486, compared to the point before) and EL increased to 75.37 ± 9.31% but the difference was not significantly different from the previous value (*p* = 0.204).

**Figure 6.**
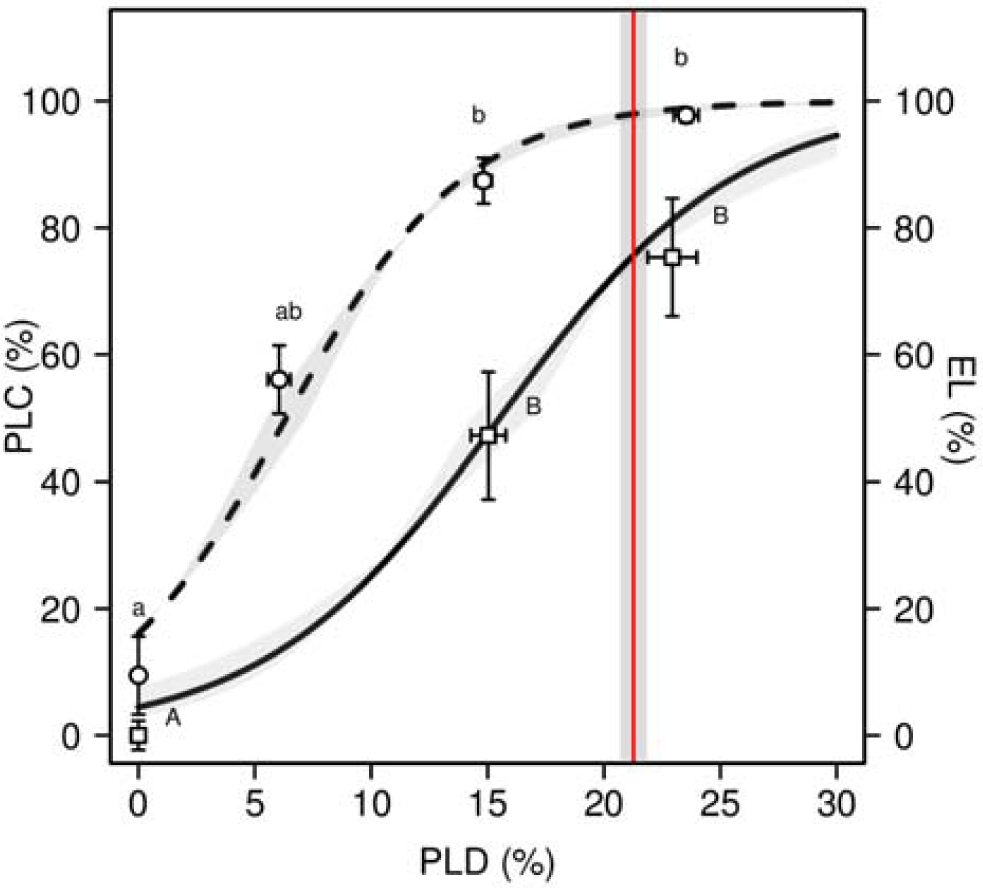
PLD-course of xylem embolism and cell lysis in *L.a.* grown under well watered condition then submitted to a drought. Data are mean values of PLC (circles) ± S.E. (n= 4 to 14) and mean values of EL (squares) ± S.E. (n = 7). Full and dashed lines represent logistic models for PLC and EL, respectively; and grey areas represent S.E. The red vertical line represents PLD_*max*_ mean value (Table 1) ± S.E. (shaded area). The different letters indicate significant differences at *p* < 0.05 *(post hoc* Dunn’s test).

Both PLC and EL were positively correlated to PLD according to a sigmoidal function (pseudo-r^2^ = 0.85 and 0.75, respectively, Fig. 6). PLD_88h_ (the PLD value for which the PLC reached 88% or hydraulically relevant PLD) was close to the PLD_50c_ (PLD value for which the EL reached 50%, or cellular relevant PLD): 14.05% and 15.52 ± 1.1 %, for PLD_88h_ and PLD_50C_, respectively. PLD_50_ (from Fig. 3) was not significantly different to PLD_50c_ (*p* = 0.070). At PLD_max_ (loss for potential recovery), PLC was near to 97.96% and EL near to 75.66%. Interestingly, loss of cell viability and recovery capacity occurred at higher PLD than the reported hydraulic failure (Barigah et al., 2013).

## Discussion

### Can the loss in branch diameter help in predicting drought-induced mortality?

Water potential measurements is classically used to measure plant water status. However, under extreme drought conditions, very few free water remains in the tissues setting a methodological limit at *ca.* −10MPa for pressure chamber and −7MPa for psychrometers. The index developed in this study (percentage loss of diameter, or PLD) is based on a length measurement and therefore can be applied in a wide range of conditions, reflecting the stress intensity the plant experiences. In this study, the diameter variations recorded along the desiccation process (from mild to extreme water stress) were non-destructive, continuously recorded with high accuracy (*ca.* ≤ 1 µm). However, the relation between PLD and water potential was not linear. The maximum measurable water potential was reached early in the dehydration while the PLD was still increasing.

The relation between stem PLRC and PLD highlighted the irreversible diameter threshold (PLRC = 100%) at PLD_max_. The PLD_max_ value was similar across branches whatever the diameter, age, growth conditions and species (21.27 ± 0.57%). Plants were able to recover as long as PLD_max_ was not reached, but not after. This index can thus be considered relevant to indicate drought-induced mortality, at least in lavender species.

This is, to our knowledge, the first reported use of LVDT dendrometers on small shrub species. The accuracy of the sensor was high enough to monitor phenological and stress events (*eg.* growth, daily fluctutations, and recovery). The mortality was assessed through the ability of the plant to recover following drought, indicated by diameter increment after rewatering and new shoots produced next growing season (Brodribb and Cochard, 2009; Barigah et al., 2013b). The sharp decrease of stem diameter during drought stress was not recovered after rewatering in dead plants, hereafter defined as the irreversible diameter threshold. We consider that the diameter monitoring on one of the main branch is relevant for the whole plant in small shrub species, as selected branches were closely linked to the trunk (see also Supplemental Table S1). After rewatering the water potential was fully restored, whatever the level of damage developed in the plant. However, damaged plants did not fully recover in diameter (PLRC > 0) indicating partial damages (Améglio et al., 2002). Indeed, in other studies irreversible stem shrinkages are related to the amount of cells injured by freezing temperature and is considered as a relevant index of frost damages (Améglio et al., 2002; Lintunen et al., 2015). PLRC appears therefore a relevant proxy for drought-induced damages. Stem PLRC would be associated to cell damages in the elastic compartments, and this is supported by the fact that PLD_50_ and PLD_50c_ were not significantly different.

Leaf PLRC was measured by weighing to test leaf viability under drought in evergreen Mediterranean species (Oppenheimer and Leshem, 1966). Recent studies used the leaf PLRC (*e.g.* John et al., 2018; Trueba et al., 2019) to help in understanding the failure of leaf function under drought and to compare species for drought tolerance. Leaf PLRC was associated to cell damages, especially by the deregulation of reactive oxygen species homeostasis (Oppenheimer and Leshem, 1966; Sharma et al., 2012), but it was not related to plant survival. Here, we used a stem PLRC not related to the full water content but to the stem diameter variations, in order to investigate mortality mechanisms under severe drought conditions, while the failure of leaf function has already occurred, as the leaves are no longer functional (Fig. 5). In addition, the used device has the advantage to monitor stem PLRC on the whole plant and not on a detached organ. This allows, among other things, to investigate the recovery phase *in planta*.

PLD_max_ in lavender species showed a total branch shrinkage of 21% on average. We did not differentiate bark and xylem component as diameter measurements integrated both compartments (over-bark). It is commonly accepted that bark, composed of several living cells, cork, parenchyma, phloem, cambia and immature xylem is extensible (Genard et al., 2001), unlike mature xylem which is considered rigid because it is lignified (Steppe et al., 2006). This subject remains controversial because some studies listed by Alméras et al., (2006) have shown that xylem can make a significant contribution to stem shrinkage and swelling, with for example 25% of the stem changes (Brough et al., 1986). In lavender species we also found that bark dehydrates more than xylem but its contraction is not enough to explain the PLD_max_. Thus, the present study clearly showed that drought-induced mortality only happens when the water supply in “elastic” tissues is fully released.

### New insights on mechanisms of woody plant mortality

In the present experiments, stems of *Lavandula angustifolia* Maillette and *Lavandula x intermedia* Grosso were relatively vulnerable to drought-induced embolism (*P*_50_ = −2.46 ± 0.14 MPa and −2.41 ± 0.09 MPa, respectively), compared to other Mediterranean species. Indeed, median *P*_50_ for this vegetation type is close to −5MPa according to Maherali et al., (2004). Two different techniques agreed in assessing *P*_50_ and highlighted the lack of differences between two lavender taxa. The clones used in the present study are the most widely used in French fields and, as often, they were only selected based on yield. We assume a large plasticity to explain high *P*_50_ values in our experiment (Awad et al., 2010; Herbette et al., 2010). The studied plants grew under favourable growth conditions whereas Lavandin plants grown under limited resources and binding conditions in the field exhibited much lower vulnerability to embolism (*P*_50_ = −5.34 ± 047 MPa; Supplemental Figure S3). Although both lavender species appeared vulnerable to cavitation, they were rather resistance to drought. The theoretical lethal water potential *P*_88_ was similar across species (−3.65 MPa and −3.76 MPa for *L.a* and *L.i*, respectively). However, rewatered plants remained able to recover, after being exposed to lower water potential than *P*_88_, as in five other tree species (Li et al 2015). Moreover, the PLD inducing 88% of PLC were lower than PLD_max_ (14.05% *vs.* 21.27% for PLD_88h_ and PLD_max_, respectively). PLD_max_ was associated to a high rate of cell lysis (Fig. 6) and so high embolism level is not sufficient to explain mortality in lavender. Besides, cavitation would contribute to the drought resistance. In grapevine, the loss of hydraulic conductivity is accompanied by an increase in hydraulic capacitance (Tyree and Yang, 1990; Hölttä et al., 2009; Vergeynst et al., 2014), suggesting that cavitation could buffered the decrease in water potential and thus help in surviving during drought.

When PLD_max_ was reached, the cambium was probably injured as well as the other elastic tissue such as parenchyma and all vessels embolized since a long time. Consequently, the growth cannot resume since there is no more meristematic cells or any parenchyma cells that can dedifferentiate. On the contrary, when PLD_max_ was not reached, the growth resumed despite a dead volume in the branches, involving an active cambium that generates new xylem vessels to recover from an episode of drought (Brodribb et al., 2010; Guadagno et al., 2017)

According to Li et al., (2015) and Guadagno et al., (2017), a threshold in the cell death (membrane injury or mortality of cambial cells) can be a point of no return associated with plant death. Maintaining the stability of cell membranes is an integral part of drought tolerance (Bajji et al., 2002). During a drought applied on poplar, the bud and cambia cells remains the last hydrated territories in a stem (Barigah et al., 2013a). The bud respiration, *i.e.* its survival, was closely associated to its water content (Barigah et al., 2013a). Here, we could not investigate bud activities but we could analyse the radial growth as a proxy of cambial activity. We thus showed that, as long as the plant can monopolize water in the elastic reservoir, cambial activity can recover. These results also showed that lavender species have a large range of capacitances from mild to severe stress, and can mobilize water reserves in sublethal drought conditions. This would be better taken into account when investigating the trait contributions in drought tolerance.

## Conclusion

The drought-induced mortality is not exclusively related to xylem hydraulic failure in lavender species, at least in these experimental conditions. The recovery is affected as soon as there is no more water storage in the stem elastic compartments. This is associated to a high level of cell damages assessed by electrolyte leakage. Continuous measurement of stem diameter variations proved to be relevant and allowed identifying a dehydration threshold for the recovery to extreme drought and related mortality. The results and tools developed in the present study would help the lavender growers to anticipate extreme drought situations that is likely to be more frequent in the future.

An important further step would be to test the genericity of these results across species. The presented work is based on the common mechanisms in woody plants and measuring PLD_max_ and PLRC in other species should be promising and relevant.

## Materials and Methods

### Plant material

Experiments were carried out on two clonal varieties of lavender (*Lavandula angustifolia* (*L.a.*) Maillette and *Lavandula x intermedia* (*L.i.*) Grosso) during the summers 2017 and 2018. Plants were produced from 1-year-old woody cuttings provided by a lavender producer in Les-granges-Gontardes (N 44° 24′ 57.24′′, E 4° 45′ 47.304′′, 100 m *a.s.l*). Plants were potted in a dedicated aromatic plant potting soil (Klasmann code 693 - medium fibrous structure) and well-watered in a greenhouse for 6 months. Homogeneous plants were selected according to their height, architecture and leaf density.

#### Experiment 1

In November 2016, selected plants were put in 7 liters pots filled with 4500 g of satured soil with water and put in a greenhouse at the horticultural school of Romans-sur-Isère (South of France; N 45° 2′ 44.16′′, E 5° 3′ 3.096′′, 250 m *a.s.l*). Pots were regularly watered to field capacity.

#### Experiment 2

In January 2018, selected plants were put in 10 liters pots filled with 5700 g of soil saturated with water and placed in a cooler environment in the greenhouse at the INRA research station of Clermont-Ferrand (N 45°77′, E 3°14′; 300 m *a.s.l.*). Temperature and relative humidity were monitored and temperature was limited to 35°C. After three months, two sets of plants were submitted to contrasted water regimes: well-watered (WW) at field capacity and water limited (WL) for each species. The WL plants were maintained at 25% to 30% field capacity, using balances and electronic valves for irrigation as described in Niez et al., (2019). Water limitation was set as the minimal water level that still allow growth (measured using Linear Variable Differential Transformer (LVDT) dendrometers).

### Drought stress and recovery

For both experiments, two successive droughts by irrigation stops (IS) were applied at the beginning of flowering of *L.i.*. The first IS was short and allowed full recovery after rehydration and the second one was more intense. Plants were rewatered successively to test the plant recovery. After the second IS, plants were considered dead when they were unable to grow and produce new shoots until the following year.

In the experiment 1, two sets of plants were treated differently: water stressed (46 individuals for *L.i* and 66 for *L.a*) and control plants (10 individuals *L.i* and 20 *L.a*). Plants were divided in blocks of ten individuals: 1 and 2 control blocks and 5 and 7 water stressed blocks for *L.i* and *L.a*, respectively. The different blocks were distributed randomly in the space. A first IS was applied in May 2017 for 4 days then plants were irrigated up to field capacity. After 4 days, a second IS was applied for various periods of time *i.e.* 10, 13, 21 and 28 days after the onset of IS. Control plants were kept watered at field capacity during the whole experiment.

In the experiment 2, 40 plants were selected. Twenty of them (10 *L.i*. and 10 *L.a.*) were grown under WW conditions, while the other 20 (10 *L.i*. and 10 *L.a.*) were grown under WL conditions. For each growing condition, 6 *L.i.* and 6 *L.a.* were water stressed, whereas 4 *L.i.* and 4 *L.a.* were control plants. A first IS (IS_1_) was applied in June 2018 for 3 days before plants were rewatered to field capacity. After 4 days, a second IS (IS_2_) was applied for various periods of time *i.e.* 8, 11, 15, 18 and 39 days for WW condition, and 22, 25, 26, 29 and 39 days for WL condition.

Eight additional *L.a.* WW from experiment 2 were used in a temperature-controlled chamber for a unique dehydration. Plants were uprooted and dehydrated until mortality, at constant temperature (25°C) and light (2 lights of 25W and 172lm). Regular measurements of electrolyte leakage, embolism and loss of diameter were made. The experiment was conducted as long as the diameter decreased (*ca.* 10 days).

### Monitoring of stem diameter variations

During all experiments, stem diameters were continuously measured using miniature displacement sensor with a friction free core sticked to the bark and Linear Variable Differential Transformers (LVDT: model DF2.5 and DF5.0; Solartron Metrology, Massy, France) connected to a data logger (CR1000, Campbell Scientific LTD, Logan, Utah, USA) or a wireless PepiPIAF system (Hydrasol, Le Plessis Robinson, France). Straight and unbranched sections of main branches longer than 5 cm were randomly chosen to mount LVDT dendrometers by a custom-made stainless Invar (alloy with minimal thermal expansion) holder. At the end of each experiment, the final stem diameter was measured using a calliper (Burg Wächter, 0.01mm accuracy) at the location of friction-free core of LVDT dendrometers measurement. The diameter time change were computed from the final diameter and the recorded variations for each individual.

In experiment 1, 20 LVDT dendrometers were mounted: 5 for each treatment and species. The branch diameter was recorded at every 30 min from March 2017 to February 2018. In the experiment 2, 32 LVDT dendrometers were mounted: 2 and 6 for control and water stressed plants, respectively for each growing condition and species. The branch diameter was recorded every 30 min from April 2018 until October 2018. In the temperature-controlled chamber, 8 LVDT dendrometers were mounted (1 per individual), at 5 min time intervals and until the end of the experiment.

Two different indexes were calculated from branch diameter variation during drought stress and recovery (Fig. 1). The Percentage Loss of Diameter (PLD) during the different IS was calculated for each plant as:

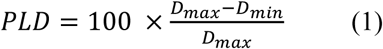

where *D*_max_ and *D*_min_ are the maximum (before the IS) and the minimal diameter (before rewatering), respectively.

When no recovery was observed after rewatering, the PLD was considered maximum (PLD_max_). When a diameter recovery was recorded following rewatering, the non-recovered diameter was calculated as the Percentage Loss of Rehydration Capacity (PLRC):

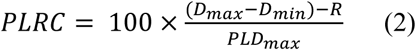

where R is the portion of D_max_ that recovered the following night after rewatering.

### Stomatal conductance

In both greenhouse experiments, stomatal conductance was measured on the abaxial surface of leaves using a porometer (AP4; Delta-T Devices Ltd, Cambridge, United Kingdom). For each kinetic point, at least 3 individual plants per condition were measured with at least 2 leaves per individual.

### Water potential

During greenhouse experiments, midday water potential was measured at the solar noon between 1:00 p.m. and 2:00 p.m. using a Scholander-type pressure chamber (PMS Instrument, Albany, OR, USA). Measurements were carried out on *ca.* 5-10 cm long stem segments from the top of a branch and bearing several leaves. For each date, measurements were performed on at least 1 individual per block (Experiment 1) or at least 3 individuals per condition (Experiment 2). For the experiment 2, at one date before the drought stress, predawn water potential was measured to compare the two growth conditions. In the temperature-controlled experiment, the water potential was measured regularly during the dehydration course. When water potential reached lower values than −9MPa, leaves were extremely dry and brittle and the measurement could not be performed.

### Xylem native embolism

During the experimentation 2, native embolism on stems was measured using a xylem embolism meter (XYL’EM, Bronkhorst, Montigny-les-Cormeilles, France). The entire inflorescence stems of about 30 cm long were collected and put in wet black plastic bags and brought them immediately to the laboratory for measurements. Segments of 2 cm were cut under water and fitted to water-filled tubing. One end of the stem segment was connected to a tank of de-gassed, filtered 10mM KCl and 1mM CaCl_2_ solution. The flux of the solution was recorded through the stem section under low pressure (60 – 90 mbar) and the initial hydraulic conductance (*K*_i_) scored. Then, the stem was perfused at least twice for 10 sec then 2 min at 1 bar until the hydraulic conductance no longer increased in order to remove air from embolized vessels and to determine the maximum conductance (*K*_max_). The percentage loss of hydraulic conductance (PLC) was determined as:

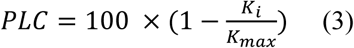

PLC was measured on water-stressed and control plants with at least 3 repetitions for each condition and date. In the temperature-controlled chamber, PLC was measured on each plant with 3 repetitions.

### Vulnerability to embolism

Vulnerability to embolism was measured on *L.a* and *L.i* using two methods. First, the vulnerability to embolism was measured using the Cavitron technique, as described by Cochard et al., (2005). Entire inflorescence stems (approx. 30 cm long) were collected from the plants in greenhouse experiment 1, put in wet black plastic bags and brought to the UMR PIAF research station of Clermont-Ferrand for measurement using Cavitron. Samples were cut and shortened to 26 cm length under water and were brought into bundles. Centrifugal force was used to establish negative pressure in the xylem and to provoke embolism, using a 28 cm diameter rotor mounted on a high-speed centrifuge (Sorvall RC5B Plus). The xylem pressure, controlled by the rotation speed of the rotor was stabilized every 0.5 MPa for 5 minutes before conductivity was measured. PLC were determined at various xylem pressure to obtain a vulnerability curve. In total, 4 and 7 vulnerability curves by Cavitron have been obtained for *L.a* and *L.i*, respectively. For the second method, the bench dehydration one, the vulnerability curves were drawn by plotting the PLC *vs.* the water potential data obtained during the drought stress in the experiment 2.

A sigmoidal function was used to fit each curve using the following equation:

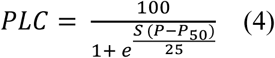

Where *S* is the slope of the curve and *P*_50_ the pressure inducing 50% of loss of conductance. The pressure inducing 88% loss of conductance (*P*_88_) were calculated as:

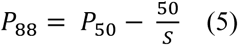

### Electrolyte leakage

Electrolyte leakage is measured to evaluate cellular damages induced by various stress (see *e.g.* Herbette et al., 2005; Charrier and Améglio, 2011). To assess drought-induced cellular damages, the electrolyte leakage (EL) was measured on branches of 8 individuals *L.a.* during the experiment performed in the chamber. Samples were cut into 5 cm long sections and immersed into 15 ml of distilled-deionized water. Vials were shaken for 24 h at +5 °C in the dark (to limit bacterial growth) on a horizontal gravity shaker (ST5, CAT, Staufen, Germany). The electric conductivity of the solution was measured (*C*_1_) at room temperature using a conductimeter (Held Meter LF340, TetraCon® 325, Weiheim, Germany). After autoclaving at +120 °C for 30 min and cooling down to room temperature, the conductivity was measured again (*C*_2_). Relative EL was calculated as:

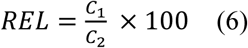

To normalize the REL, 10 control samples and 10 samples were frozen at −80°C and used to have a reference for 0 (REL_WW_) and 100% cellular damages (REL_-80_), respectively. An index of damages *I*_Dam_ was computed as:

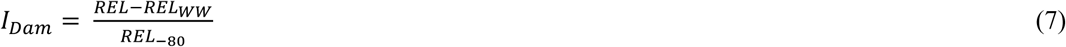

## Statistical analysis

Statistical analyses were performed using the RStudio software (under R core version 3.4.3, R Development Core Team, 2017). Kruskal-Wallis test was used for the group comparisons between conditions for PLD_max_, PLC and *I*_dam_, followed by a *post hoc* Dunn’s test when effects were significant. Student test was used to compare vulnerability curves between species. The nls function was used to fit the relation between PLRC, PLC, EL and PLD. All statistical analyses were based on a 0.05 significance level.

## Supplemental Data

Supplemental Figure S1. All the dynamics of the stem diameter variations versus time.

Supplemental Figure S2. Diameter and temperature variations versus time in one *L.i.* plant alive and one *L.i.* plant dead after a rehydration.

Supplemental Table S1. PLC measured all along the shoot from *L.a.* and *L.i.* plants, on sections of 2 cm.

Supplemental Figure S3. Vulnerability to cavitation of *L.i.* plants grown in the field with no watering.

## Acknowledgements

The authors thank J. Roustand for providing material plants. P. Danelon, C. Baconnier and V. Fazekas from Lycée horticole Terre d’Horizon for plant care and handling during the first greenhouse experiment. They thank P. Chaleil and A. Faure for plant care in the second greenhouse experiment. They thank R. Souchal, P. Conchon, J. Cartailler and C. Serre for irrigation system and LVDT preparation. They thank F. Sabin for his help in PLC and EL measurements. Finally, the authors acknowledge Crieppam, Chambre d’agriculture Drôme and Vaucluse, The French Essential Oils Interprofessional Association and Iteipmai for initiating and contributing to the project.

